# DIRT/3D: 3D root phenotyping for field grown maize (*Zea mays*)

**DOI:** 10.1101/2020.06.30.180059

**Authors:** Suxing Liu, Carlos Sherard Barrow, Meredith Hanlon, Jonathan P. Lynch, Alexander Bucksch

## Abstract

The development of crops with deeper roots holds substantial promise to mitigate the consequences of climate change. Deeper roots are an essential factor to improve water uptake as a way to enhance crop resilience to drought, to increase nitrogen capture, to reduce fertilizer inputs and, to increase carbon sequestration from the atmosphere to improve soil organic fertility. A major bottleneck to achieving these improvements is high-throughput phenotyping to quantify root phenotypes of field-grown roots. We address this bottleneck with DIRT/3D, a newly developed image-based 3D root phenotyping platform, which measures 18 architecture traits from mature field-grown maize root crowns excavated with the Shovelomics technique. DIRT/3D reliably computed all 18 traits, including distance between whorls and the number, angles, and diameters of nodal roots, on a test panel of 12 contrasting maize genotypes. The computed results were validated through comparison with manual measurements. Overall, we observed a coefficient of determination of *r*^2^>0.84 and a high broad-sense heritability of 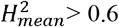 for all but one trait. The average values of the 18 traits and a newly developed descriptor to characterize a complete root architecture distinguished all genotypes. DIRT/3D is a step towards automated quantification of highly occluded maize root crowns. Therefore, DIRT/3D supports breeders and root biologists in improving carbon sequestration and food security in the face of the adverse effects of climate change.

## Introduction

Evaluating the information encoded in the shape of a plant as a response to environments is essential to understand the function of plant organs (Bucksch, 2011). In particular, roots exhibit shape diversity that is measurable as variation in rooting angles, numbers of roots per type, length, or diameter of roots within a root crown (Lynch and Brown, 2012). An understanding of variation in root crown architecture facilitates breeding for favorable root characteristics to improve yield in suboptimal conditions, including those resulting from climate change (Lynch, 2013). Improving root phenotypes through crop breeding and management holds promise for improved food security in developing nations, where drought, low soil fertility and biotic constraints to root function are primary causes of low yields, and also for reducing the environmental impacts of intensive agriculture by reducing the need for intensive fertilization and irrigation (Lynch, 2019). Root traits also offer opportunities to improve carbon sequestration (Paustian et al., 2016). These challenges demand efforts across a range of disciplines, from mathematics over computer science to plant biology and applied fields like plant breeding and agronomy (Bucksch et al., 2017; Bucksch et al., 2017).

A major interdisciplinary challenge in root biology is the development of deeper rooting crop varieties. Deeper roots promise a three-fold impact: they improve drought resilience, lower fertilizer input, and decrease atmospheric carbon. Deeper roots improve drought resilience to stronger and more frequently occurring droughts (Ault, 2020) by tapping into water in deep soil domains (Lynch and Wojciechowski, 2015). Nitrogen capture increases when roots grow deeper because nitrogen diffuses into and accumulates in deeper soil layers (Lynch, 2019). Deeper rooting crops increase carbon sequestration (Smith et al., 2007) mostly by depositing more organic residues in soils, thereby replenishing carbon after harvest (Paustian et al., 1997). An important tool in breeding deeper roots is the large-scale automated evaluation of root traits in the highly occluded root crown architecture (Topp et al., 2016). Maize (*Zea mays*) in particular, with over 700 Mt of maize production world-wide (Ranum et al., 2014), is a prime target for improving rooting depth. However, measuring important root traits for deeper rooting (Lynch and Wojciechowski, 2015) is hampered by the availability of advanced root phenotyping methods on the field-scale (Kuijken et al., 2015). Therefore, the research community has pushed for the development of a root phenotyping system that operates under field conditions (Paez-Garcia et al., 2015).

### Root phenotyping in the field remains a significant challenge for root biology

The majority of existing phenotyping methods to evaluate root architecture non-destructively emerged from laboratory settings. These methods range from fully automatic (Galkovskyi et al., 2012) to manually assisted (Lobet et al., 2011) and consider a variety of growth systems like gel cylinders (Iyer-Pascuzzi et al., 2010), rhizotrons (Nagel et al., 2012; Rellán-Álvarez et al., 2015; Bontpart et al., 2020), mesocosms (Nagel et al., 2012) and germination paper (Falk et al., 2020). Root phenotyping under lab conditions necessitates the use of constrained growth containers (Poorter et al., 2012; Bourgault et al., 2017), artificial growth media (Oliva and Dunand, 2007), and environments that potentially alter root system architecture (de Dorlodot, Bertin et al. 2005, Draye, Thaon et al. 2018). Therefore, it is essential to translate phenotyping experiments from the lab into repeatable field experiments (Zhu et al., 2011; Bucksch et al., 2014; Poorter et al., 2016).

In contrast, root phenotyping in the field is currently either invasive or destructive. Invasive procedures record small sections of the root with minirhizotron cameras placed in the soil (Gray et al., 2013; Yu et al., 2019). These procedures are incapable of recording the full root system. Therefore, investigating root system architecture in the field relies on the destructive excavation of the root crown developed in a real target environment. Shovelomics is the standard field-ready protocol to excavate root crowns in the field (Trachsel et al., 2011). It was initially developed for maize and has undergone a constant refinement by the root research community (Zheng et al., 2020). Other crops, including common bean (*Phaseolus vulgaris*) and cowpea (*Vigna unguiculata*) (Burridge et al., 2016), wheat (*T. aestivum*) (Slack et al., 2018; York et al., 2018), rapeseed (*Brassica napus*) (Arifuzzaman et al., 2019) and cassava (*Manihot esculenta*) (Kengkanna et al., 2019) have specialized shovelomics protocols in place. However, the manual excavation of the root crown, followed by visual scoring and manual trait measurement, is difficult and subjective to the researcher.

### Digital imaging of root traits in 2D enabled researcher independent large-scale analysis of field data

In response, software to measure root traits in simple digital images became available. Approaches to record root traits in the field use different methods in terms of software platforms and imaging setups. According to the software catalogue “The quantitative plant” (Lobet, 2020), DIRT is the only online platform (Das et al., 2015). The DIRT platform provides image processing and storage for over 750 root researchers following an easy to reproduce imaging protocol. For imaging, DIRT needs a tripod, a consumer camera, and a black background with a white circle. Just recently, DIRT enabled projects associated with root architecture and micronutrient content (Busener et al., 2020) and translated traits from the lab to the field (Salungyu et al., 2020). More sophisticated and expensive imaging setups use specialized tents (Colombi et al., 2015) and carefully designed imaging boxes (Grift et al., 2011; Seethepalli et al., 2019), along with computationally simple traits, for use on personal computers. The user can, therefore, choose a system that suits the project needs. Systems generally vary in the number of instruments, tools, and samples transported between the lab and field site, as well as the cost of the imaging setup and hardware requirements. However, all these systems share a single obstacle: Resolving the highly occluded branches of a dense 3D root crown. The occlusion challenge arises when the 2D image projection of the 3D branching structure ‘hides’ information of branching locations. Hence, the branching information is unrecoverable and lost (Bray and Topp, 2018).

### Unavailability of digital imaging of root traits in 3D hampers new breakthroughs in root biology

3D approaches are capable of resolving even highly occluded branching structures (Bucksch, 2014). Gel systems were among the first methods to measure fully resolved 3D root systems of younger roots (Clark et al., 2011; Topp et al., 2013) and to capture some of their growth dynamics (Symonova et al., 2015). The bottleneck of imaging and measuring older root systems in constrained growth containers filled with soil, however, remained. Therefore, X-Ray computed tomography (CT) became a widely used tool to phenotype roots in pots filled with soil and soil-like substrates (Pfeifer et al., 2015; Gerth et al., 2021). These lab developments revealed new characteristics in maize root development (Jiang et al., 2019). The X-ray CT approach is, in its applications, comparable to magnetic resonance imaging (MRI) (Metzner et al., 2015). MRI also provides a 3D model of the root (van Dusschoten et al., 2016) and can be used for time-lapse imaging of growth processes (Jahnke et al., 2009). The benefits of both X-Ray CT and MRI are substantial and subject to scientific discussion (Fischer et al., 2016). Similarly, neutron radiography can record root system architecture in soil filled growth containers (Moradi et al., 2011) to quantify water uptake of different root classes in maize (Ahmed et al., 2018). However, X-Ray CT, MRI and neutron radiography do not meet the needs for large-scale field studies: Firstly, both technologies restrict plant growth to the size of a given growth container. The restriction to smaller sizes is proportional to higher achievable resolution. Therefore, it is common to observe an immature “pot phenotype” instead of a relevant phenotype grown in field soils (Poorter et al., 2012; Bourgault et al., 2017). Secondly, both methods can take about 30 minutes or more to collect root imaging data in soil. Extremes of several days are reported for X-Ray CT systems to achieve the resolution of root hairs (Keyes et al., 2013; Sozzani et al., 2014). An additional constraint is the cost of constructing, operating, and staffing such facilities, few of which are devoted to root studies.

In response to these phenotyping limitations, we developed DIRT/3D as an automatic 3D root phenotyping system for excavated root crowns grown in agricultural fields. Our approach consists of a newly developed 3D root scanner and root analysis software. The 3D root scanner captures image data of one excavated maize root in about five minutes. Our software uses the image data to produce a colored 3D point cloud model and to compute 16 root traits. The computed traits measure individual roots and also characterize the complete root crown. Individual root traits include number, angle, and diameters of youngest and 2^nd^ youngest nodal roots. Traits like eccentricity or the distance between whorls characterize the root crown. The computed traits are known to be relevant and reported frequently in literature as manual measurements (Saengwilai et al., 2014; Zhan et al., 2019). We also introduce a new 3D whole root descriptor that encodes the arrangement of roots within the root crown with improved distinction compared to 2D whole root descriptors.

## Results

### DIRT/3D enables automatic measurement of 3D root traits for field-grown maize

We developed DIRT/3D (Digital Imaging of Root Traits in 3D) system to phenotype excavated root crowns of maize (Figure 1). The system includes a 3D root scanner and a suite of parameter-free software that reconstructs field-grown maize roots as a 3D point cloud model. The software also contains algorithms to compute 18 root traits.

**Figure 1:**
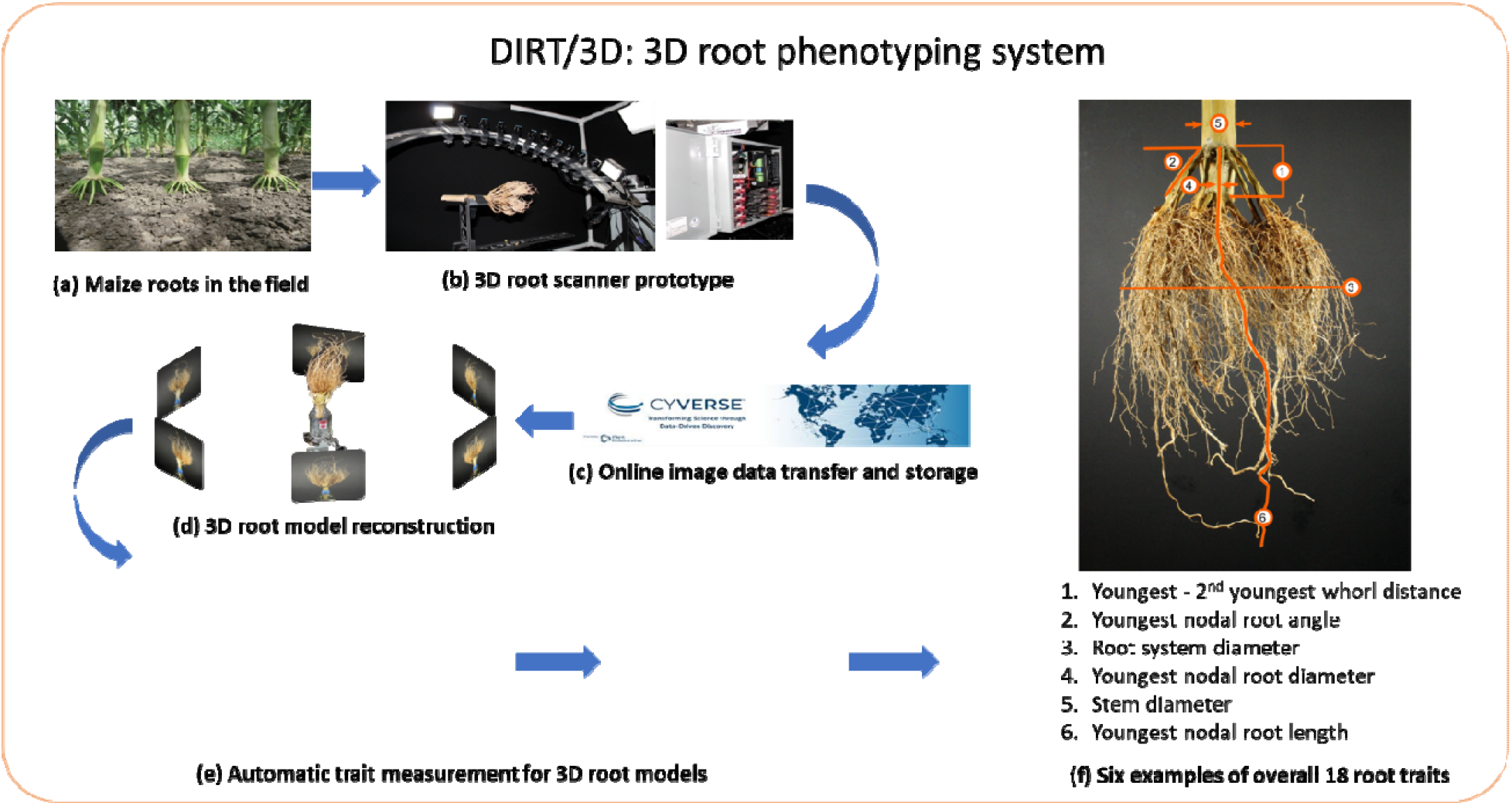
Schematic overview of DIRT/3D system. Field grown roots (a) excavated with the Shovelomics protocol (Trachsel et al., 2011) are placed in the 3D root scanner (b). The scanner, with ten synchronized industrial cameras mounted on a curved frame, acquires about 2000 images of the root. The images are transferred to and stored in the CyVerse Data Store (Merchant et al., 2016) (c). The 3D reconstruction is computed with DIRT/3D s structure-from-motion software (d) and yields the resulting 3D root model (e). Overall, the analysis software calculates 18 root traits from the 3D point cloud of the root crown. The image in (f) shows examples for the trait classes angle, diameter, and length. All developed hardware designs are open and software methods are open source. Executable are available as a Singularity or Docker container (Kurtzer et al., 2017).

The 3D root scanner (Figure 2) utilizes ten industrial cameras mounted on a rotating curved frame (Figure 2a) to capture images from all sides of the maize root (Supplementary Material 1). Scanning of one maize root completes in five minutes. After obtaining the image data, we reconstruct a 3D point cloud of the root crown. By analyzing thin level sets of the 3D point cloud, DIRT/3D revealed traits behind multiple layers of occlusions (Detailed pipeline in Supplementary Material 2). For example, DIRT/3D measures the distance between the root forming whorls and the number of nodal roots at young nodes. DIRT/3D also tracks individual roots within the root crown, starting from the stem down to the emerging lateral in the root crown. Each individually tracked root enables the measurement of numbers, angles, and diameters at the individual root level (Figure 1f).

**Figure 2:**
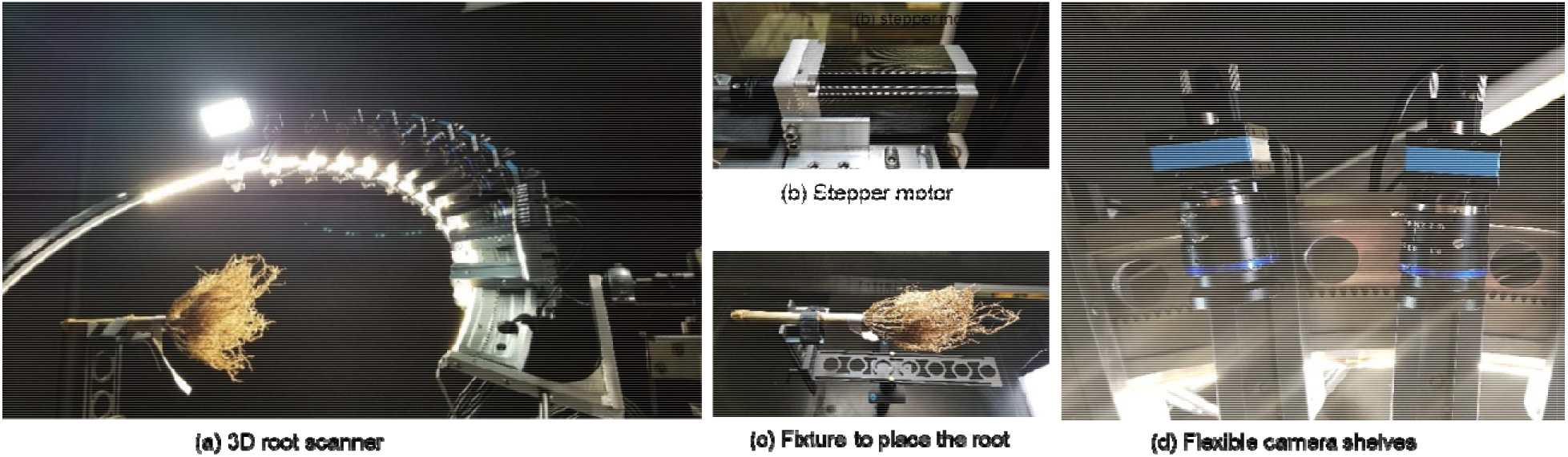
3D root scanner prototype. (a) 3D root scanner captures images of an excavated maize root grown under field conditions. (b) The stepper motor rotates the curved metal frame with the mounted cameras around the root. (c) The fixture keeps the root in place during scanning. (d) The adjustable camera shelves allow for the free positioning of each camera. The CAD drawings of the 3D root scanner are available in Supplementary Material 3.

We used a panel of 12 maize genotypes with 5-10 replicates per genotype to validate the DIRT/3D pipeline. For our validation trial, the 3D root scanner captured images at pan intervals of 1 degree and tilt intervals of 10 degrees. Figure 3 shows a visual comparison of the captured root architectural variation between the genotypes.

**Figure 3:**
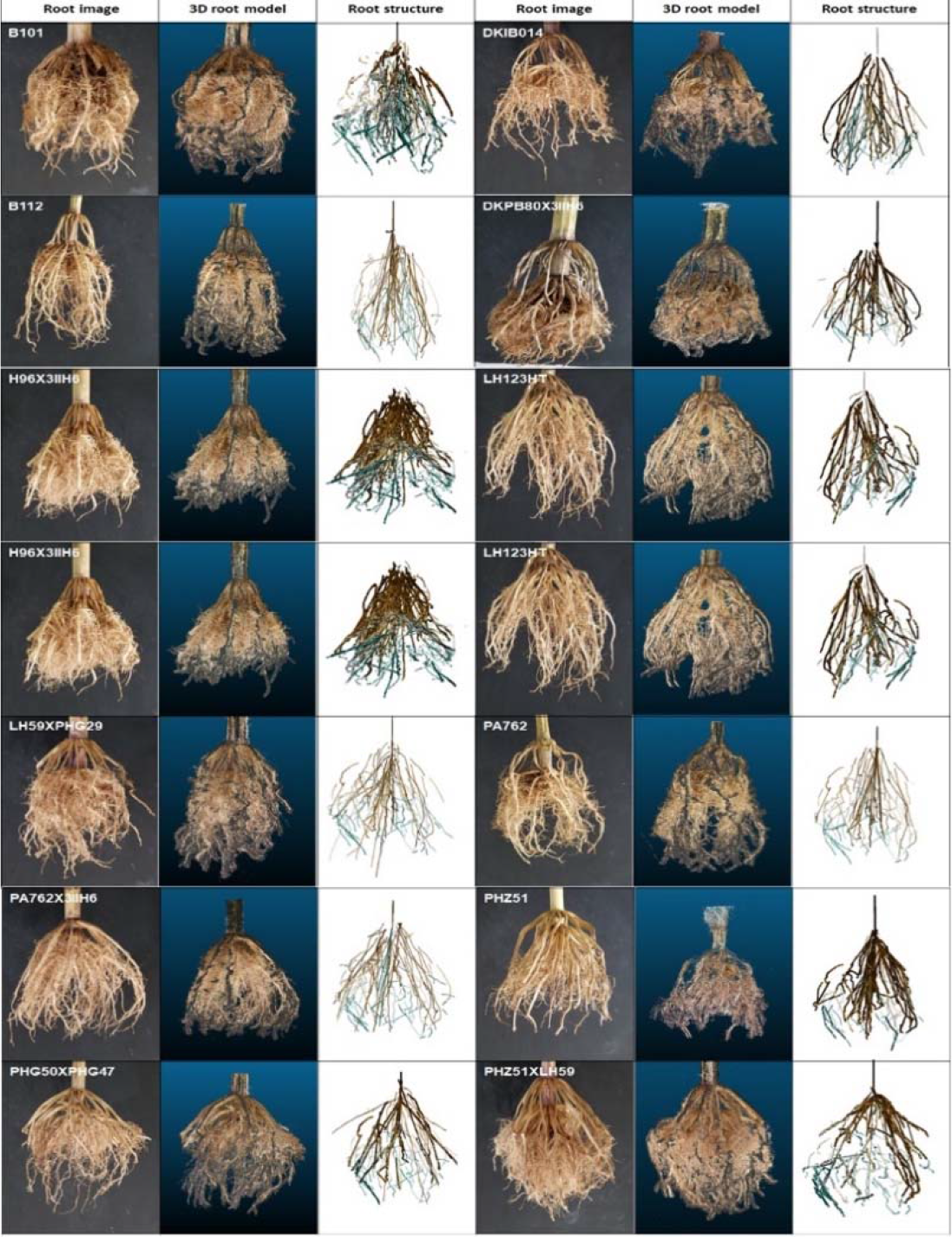
The automatic DIRT/3D pipeline generates a detailed 3D point cloud of excavated roots. Examples show excavated maize root crowns, their 3D point cloud models and structure graphs from the test panel of 12 genotypes. Visual comparison of the 2D views of real roots and their respective 3D root models shows that root architecture, along with the color, is reconstructed (Supplementary Material 4). All 3D models used in the paper are available as .ply file in Supplementary Material 5.

### Level set scanning enables extraction of traits from 3D root models

We developed a new method to perform a top-down level set scan of the 3D root model to compute 3D root traits. For a vertically aligned model, we slice the 3D root model from top to bottom at consecutive depth levels (Figure 4). The number slices represent the constant imaging volume of the scanner and therefore, vary by root crown size. Two benefits result from the fixed scanning volume. First, the transformation to mm is constant, and second, the optimal slice thickness can be determined experimentally for all roots. Therefore, all parameters are constants in the algorithm. Here, a level set image is the commonly used vertical 2D projection of each slice onto a plane (Bucksch, 2011; Mairhofer et al., 2012; Cochard et al., 2015; Dinas et al., 2015; Hyun et al., 2016) representing the sequential distribution of roots into deeper soil levels (Figure 4 (b)-(e)).

**Figure 4:**
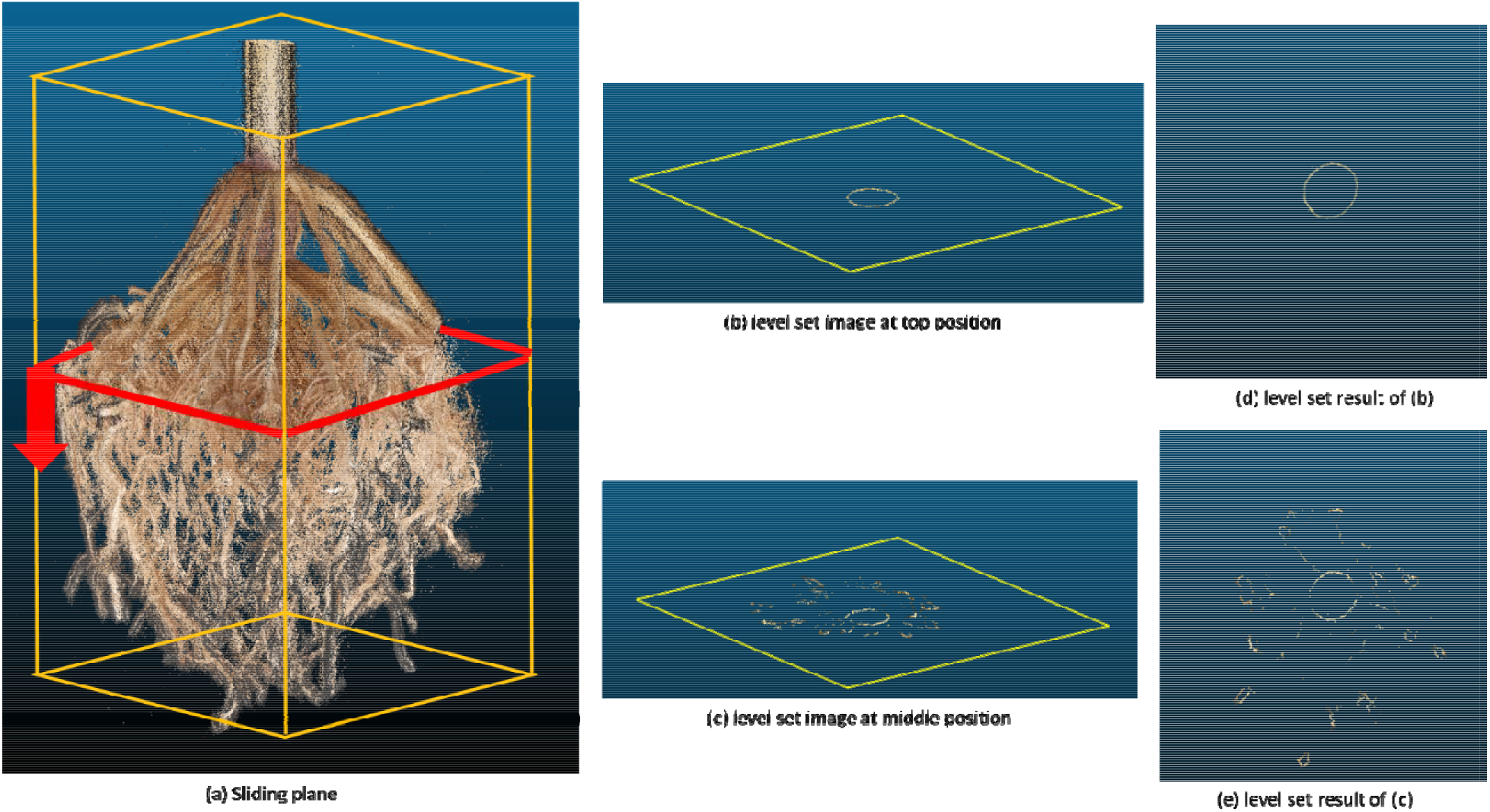
The level set image sequence for the estimation of root traits. (a) A sliding plane scans the 3D root model from top to bottom to acquire a level set image sequence. The information content per level set image varies with depth and generally encodes the points at a pre-defined distance to the sliding plane. For example, at the top (b and d), only information about the stem appears in the level set image. At a middle position (c and e), individual roots are visible as additional circles.

Ideally, each root is a closed circle in the level set plane. However, some roots are under-sampled or affected by noise such that the contours of some roots are disconnected. A video sequence of all level set images would result in a flickering effect. Therefore, we use a phase-based frame interpolation technique (Meyer et al., 2015) to smooth the level set image sequence. This method estimates transition frames between the level set images, which is equal to an up-sampling process of the 3D point cloud (Waki, 2016). Insufficient sampled locations of the root models are interpolated, which results in a smooth connection of formerly disconnected contours on level set images (Zhao et al., 2018). A comparison of the original and smoothened sequence of level set images shows the increased density of the 3D root model (Supplementary Material 6).

### The active contour snake model identifies individual roots per level set image

The image sequence of the smoothened level sets is the key to compute the location and size parameters of individual roots. Applying the active contour snake model (Kass et al., 1988) to each level set image results in a curve that circumscribes each individual root in the level set image (Mugerwa et al., 2019). Each curve contracts and moves towards the closed boundaries of an individual root by minimizing a partial differential equation, where image boundaries represent a low energy state for the active contour. The partial differential equations formulate a trade-off between an internal and external energy term. The internal energy describes the continuity and smoothness of the contour to controls for curve deformations, and an external energy that describes how well the contour fits the individual root (Zhao et al., 2018).

Our algorithm initializes a circle around each individual root in each level set as an initial input to the minimization of the active contour snake (Supplementary Material 8). During the iterative minimization of the energy function, we use a periodic boundary condition to enforce a closed curve. The resulting closed boundary curves represent first estimates of individual roots and used as input to compute a binary mask for each level set image sequence using Otsu’s binarization method (Moghaddam and Cheriet, 2012). We adopt the connected components labeling method to distinguish and label each closed-boundary object, representing individual roots (Playne and Hawick, 2018). The result of connected components labeling is a multiple segmentation of individual roots represented by colored components (Figure 5).

**Figure 5:**
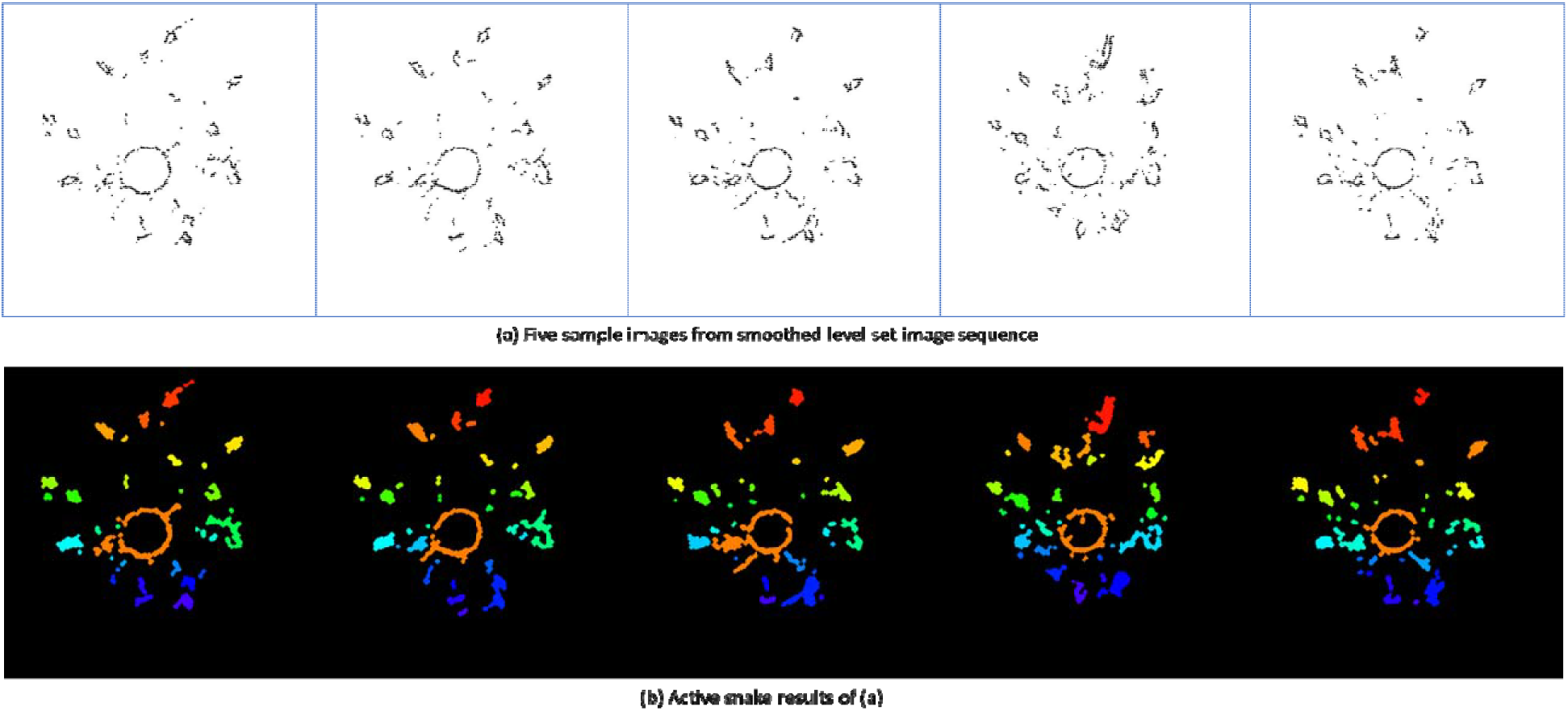
Active contour snake method identifies individual roots by analyzing connected components. (a) shows five sample images from the smoothened level set images. (b) shows the corresponding results from the active contour snake method. Each extracted individual root is color-coded with connected component labeling. The active contour snake model detects individual roots in each level set image. A video showing the active contour snake evolving process is available in Supplementary Material 7.

However, maize roots intertangle and adhere to each other, resulting in a complex system. In an image of the level set sequence, the entanglement will be visible as one connected component instead of two distinct components. We use the watershed segmentation to segment the overlapping root (Supplementary Material 9). The idea behind the watershed algorithm is to interpret grey values in the image as a local topography or elevation. The algorithm uses pre-computed local minima to flood basins around them. The algorithm terminates the flooding of a basin when the watershed lines of two basins meet. The Euclidean Distance Transform (EDT) of the image allows for direct detection of the local minima (Fabbri et al., 2008). In that way, watersheds assign each pixel to a unique component and allows the distinction entangled roots per level set image (Roshanian et al., 2016).

### A combination of Kalman filters and the Hungarian algorithm tracks individual roots

We developed an individual root tracking method by adopting a combination of Kalman filters and the Hungarian algorithm (Sahbani and Adiprawita, 2016). The algorithm detects individual roots for consecutive level set images. Once individual roots are detected, the Hungarian algorithm matches the corresponding individual roots across the level set images. To improve the speed of the Hungarian algorithm, we use a Kalman filter to predict matching individual roots in consecutive level set images (Figure 6). Behind the scenes, the tracking algorithm builds a mathematical model of expected depth development of the root. In doing so, the algorithm uses the current position, relative speed, and acceleration of individual roots to predict their location in the following level set image. As a result, we obtain an initial root structure directly from the point cloud (see Figure 3 for examples of all 12 genotypes). An animation and video showing the individual root tracking process are available in Supplementary Material 10.

**Figure 6:**
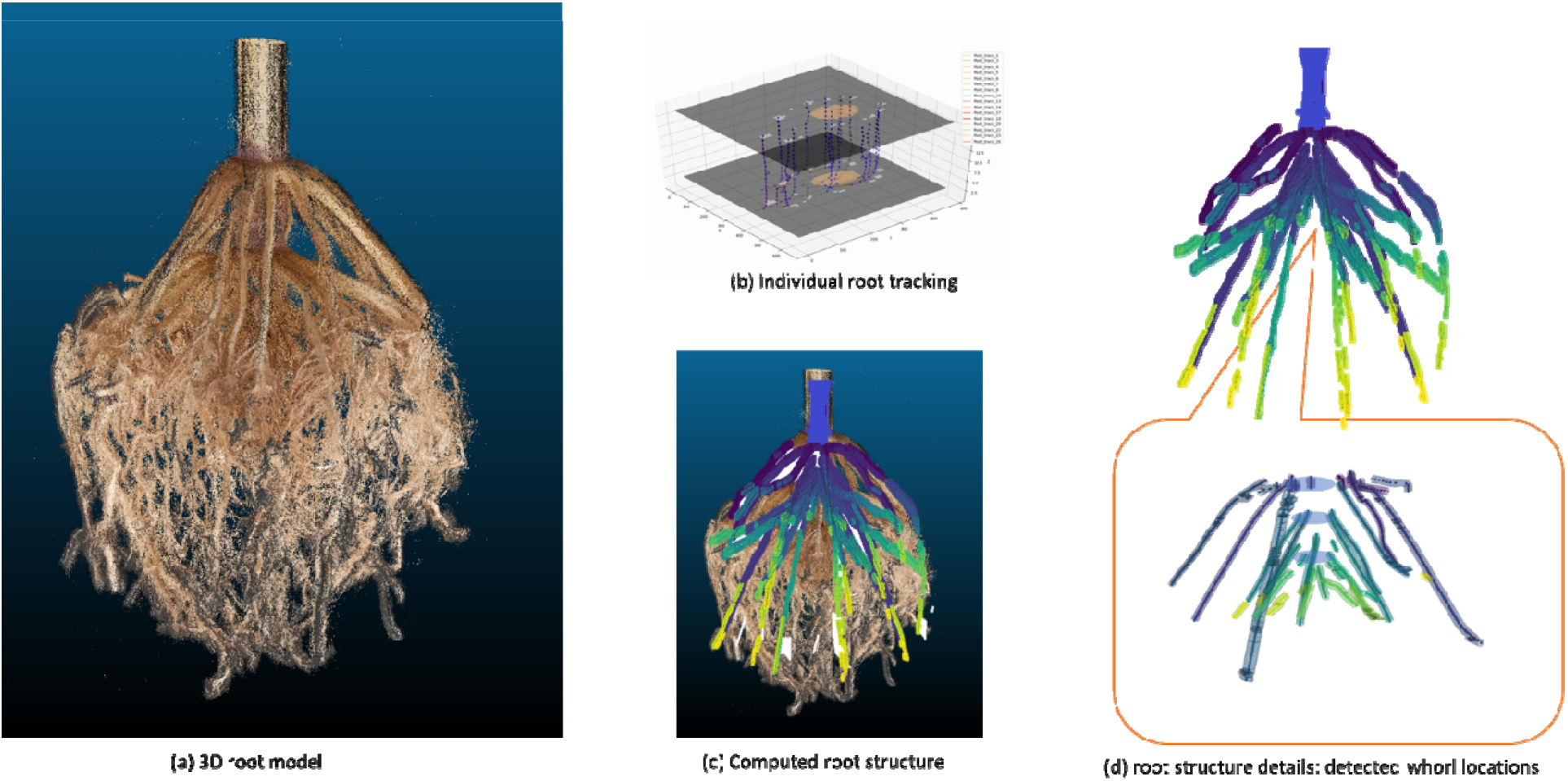
A combination of Kalman filters and the Hungarian algorithm tracks individual roots in the root crown. (a) 3D model of a field-grown maize root generated by the DIRT/3D reconstruction. (b) Visualization of the tracking process of individual roots from level set images. Two level set images are visualized with 50% transparency to show the tracking of individual roots in 3D space. (c) 3D visualization of the root structure graph consisting of all tracked trajectories of resolved nodal roots. (d) The structure graph data includes the detected whorls and the corresponding nodal roots. Individual roots are colored by depth relative to the root crown.

### Trace connection and back tracking to improve the computed root structure

Under-sampled regions within the point cloud can result from left over soil that blocks the view into the root crown. Technically, this may lead to unconnected roots in the 3D model of the root crown. As a solution to the disconnection problem, we describe each root segment by its curvature and Euclidean distance between every pair of adjacent roots and root segments. If two close-by root parts have similar curvature value, we connect them by interpolating a curved connection between both segments. We accept two connected segments as a valid solution if the newly connected segment does not deviate from the interpolated curve. Once we connected all root segments that fit the same curve, we adopt the Iterative Back-Tracking method (Liu et al., 2018) to connect remaining root segments to the root structure either as continuous curve or as a new branching root.

During the sequential processing of all level set images, we calculate the diameters of the minimal bounding circle that covers all points in all level set images in a 2D projection. Table 1 lists all 18 root architecture traits that DIRT/3D computes.

**Table 1:**
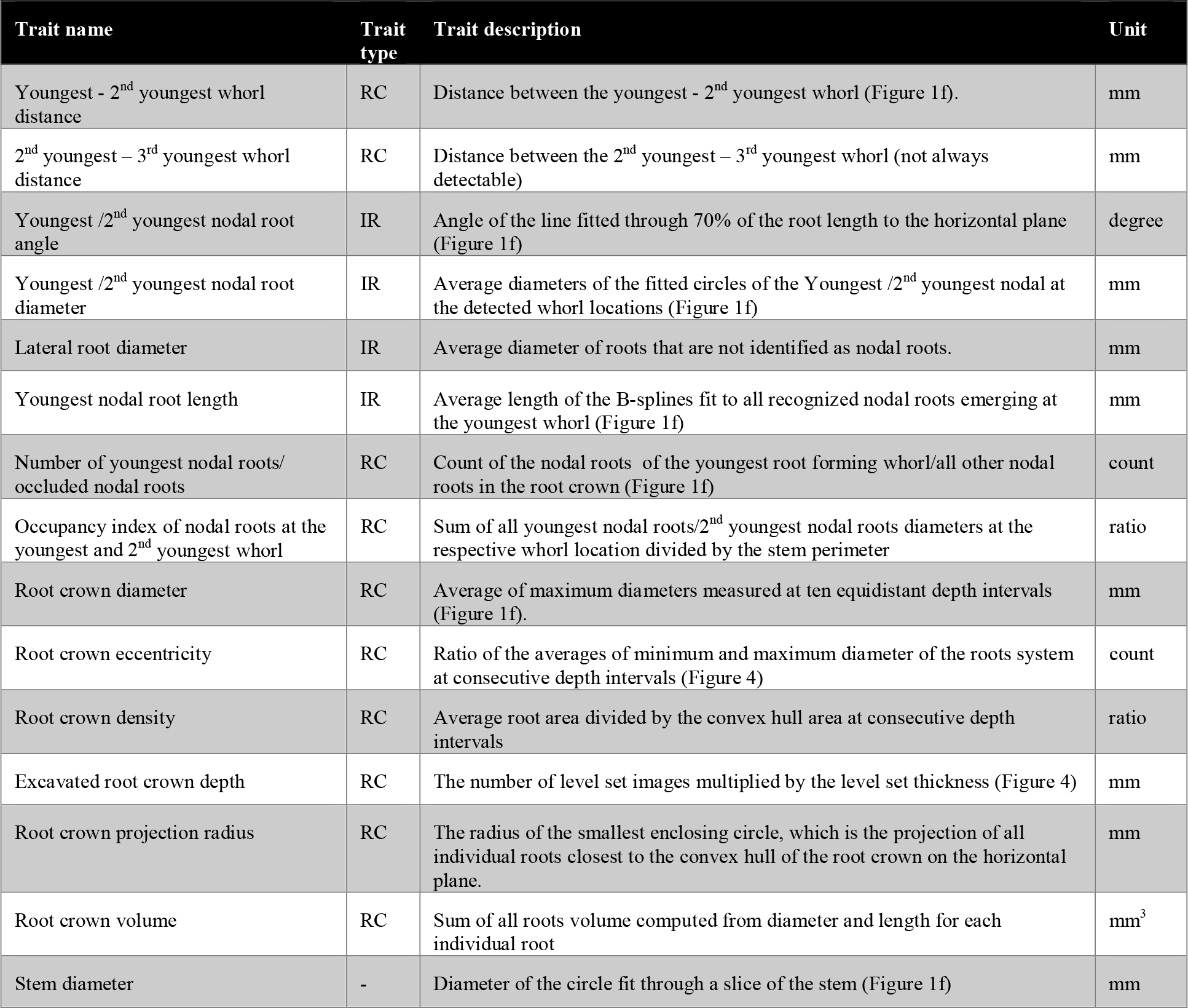
Description of DIRT/3D traits. Traits describe either a root crown (RC) characteristic or measure an individual root (IR) within the root crown.

### 3D root traits correlate at individual root and root crown level

To test the accuracy and precision of DIRT/3D, we correlated the trait values measured automatically in the 3D point clouds with manually measured traits of the root crown. We validated manually measurable traits such as root crown diameter, whorl distances, number of nodal roots at certain whorls, nodal root angle and root dry biomass with a precision scale to correlate it with root volume. The correlation analysis of the each of all validated traits showed *r*^2^>0.84 and *P* < 0.001 (Figure 7 a-d are selected examples among 10 traits validations in Supplementary Material 11). The results for the 2nd youngest – 3rd youngest whorl distance (Supplementary Material 11) indicate that at least 2 mm of whorl distance is needed to identify whorls with our methods. The minimal whorl distance sets a technical limit to distinguish the earliest whorls in the root crown.

**Figure 7:**
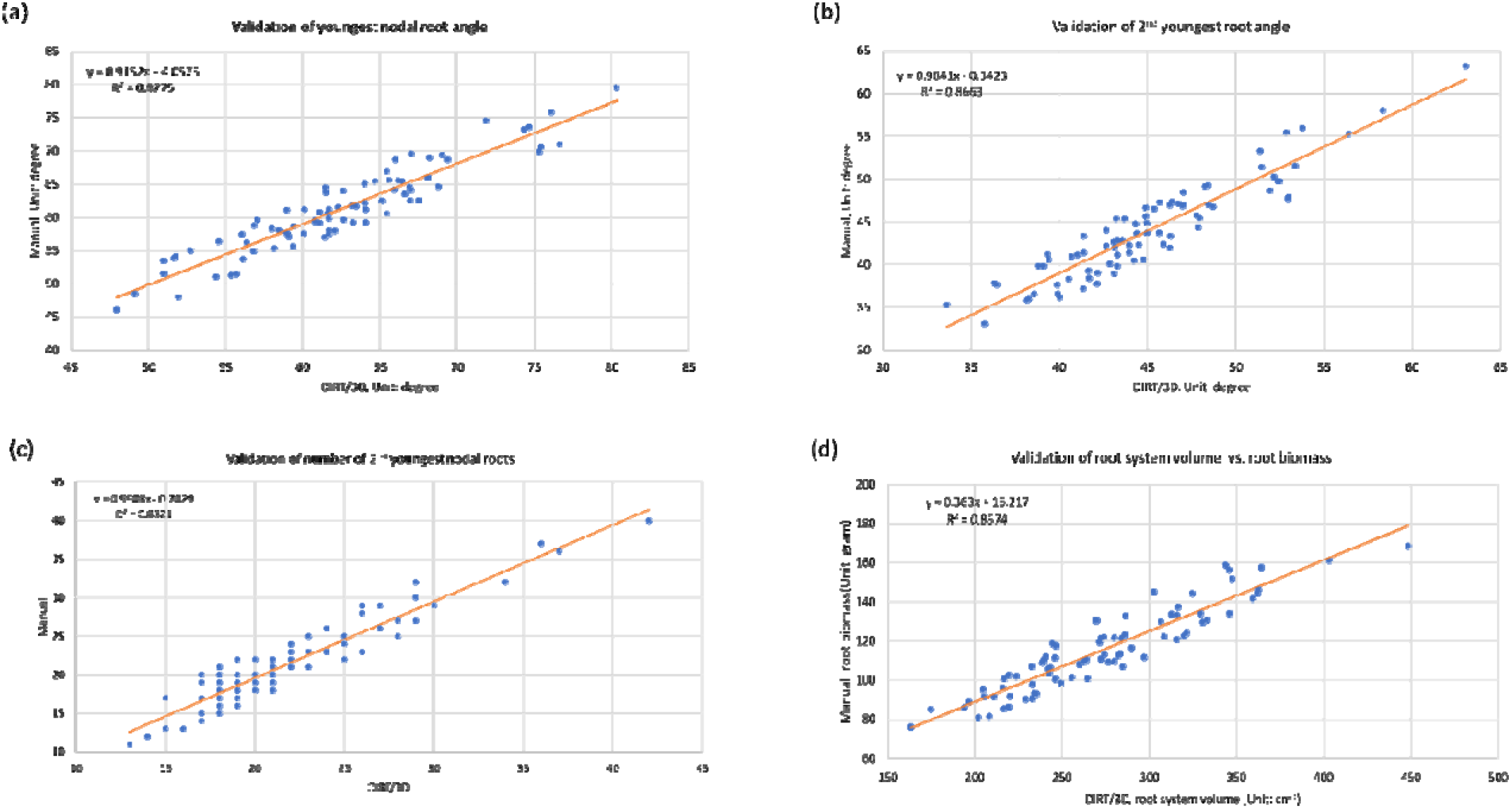
Correlation of automatic and manual trait measurements. All measured traits resulted in correlations of r^2^>0.84 (See Supplementary Material 11). The Figure above shows the examples of (a) the minimally occluded nodal root angle of the youngest whorl; (b) nodal root angle of the second youngest whorl which is occluded by the nodal roots of youngest whorl; (c) the number of nodal roots nested inside the youngest whorl; (d) root crown volume extracted with the root tracing algorithm that generates the whole root descriptor correlated against manually weighed dry biomass.

### Broad sense heritability suggests high repeatability of the observed root trait values

Broad-sense heritability, for all traits (Figure 8) is computed as the ratio of total genetic variance to total phenotypic variance (Falconer, 1989) to demonstrate the repeatability of the initial fields trial. For quantitative plant traits, the broad-sense heritability across multiple varieties eliminates the time-consuming steps of hybridization and population development for determining. We observed a broad-sense heritability for all traits except youngest nodal root length, which indicates a strong genetic basis for these traits. Nine of the computed traits resulted in (Figure 8), which indicates that the calculated traits show minimal variation within genotypes sampled with the Shovelomics method. Note that the crow-crown whorl distance could not be included into the heritability calculation because it is not always detectable at the resolution of our system.

**Figure 8:**
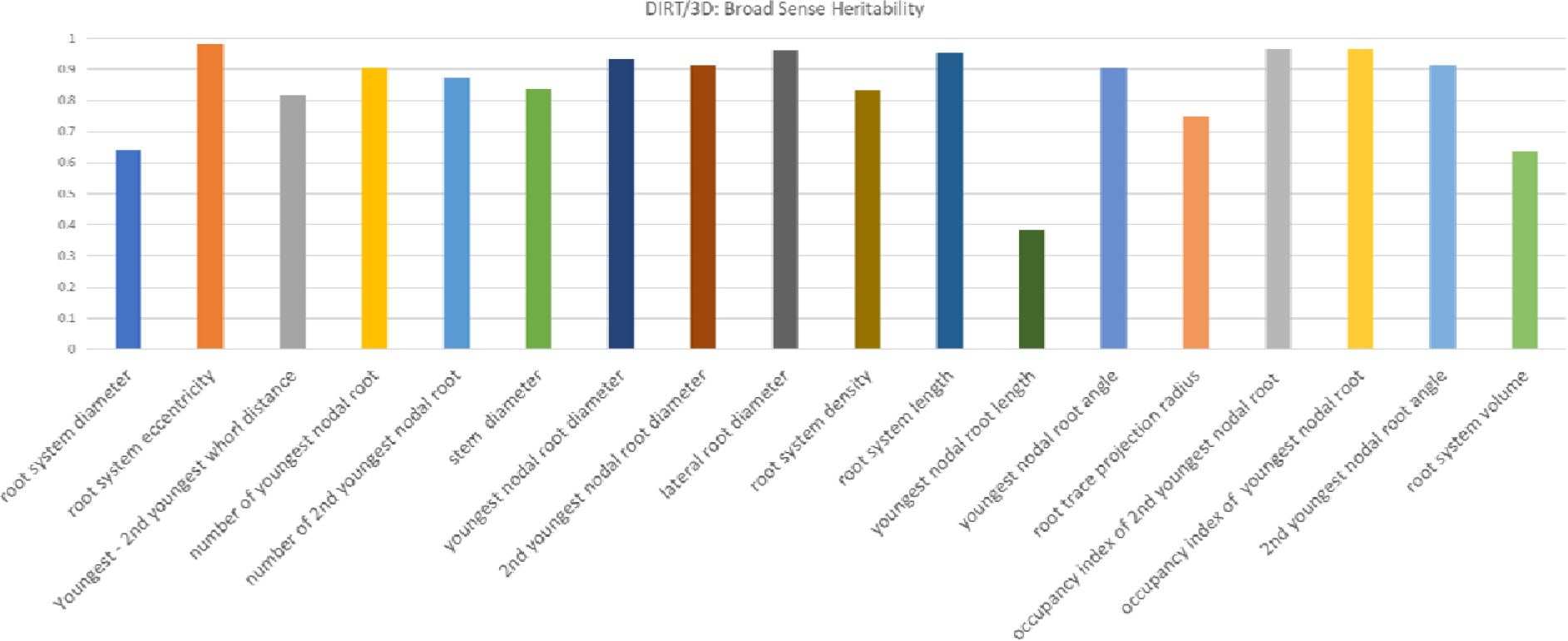
Broad sense heritability for 18 computed traits. Phenotypes vary between the individuals because of both environmental factors, the genes that control traits, as well as various interactions between genes and environmental factors. We computed broad-sense heritability for all 3D traits in Table 1. All but one trait suggests a moderately to strong genetic basis to explain the observed inter-genotypic variation with > 0.6.

### 3D Root crown traits show consistent results compared to DIRT 2D traits

We computed comparable root crown traits with DIRT/2D from images of the same roots used for the 3D analysis. Overall, four traits are comparable between DIRT/3D and DIRT/2D and 17 traits are only available with DIRT/3D. Three of the four traits are aggregate traits that vary dependent on the phenes composing them (Rangarajan and Lynch, 2021). Our comparison is possible because the 2D and 3D images represent in our case the same individual root. The four compared traits resulted in highly similar correlations with the manually determined ground truth and in similarly high broad sense heritability (Table 2). The heritability of all DIRT/2D traits can be found in Supplemental Material 16.

**Table 2:** Comparison of DIRT/3D and DIRT/2D traits for the data set of 100 maize roots used in the 3D analysis before.

| Trait name DIRT/3D vs DIRT/2D | R<sup>2</sup> DIRT/3D | R<sup>2</sup> DIRT/2D | H<sup>2</sup> DIRT/3D | H<sup>2</sup> DIRT/2D |
| --- | --- | --- | --- | --- |
| Youngest nodal root angle/Top angle | 0.8775 | 0.8377 | 0.9056 | 0.8920 |
| Root crown diameter/Median width | 0.8878 | 0.8756 | 0.6415 | 0.6125 |
| Excavated root crown depth /Root system depth | 0.9126 | 0.9153 | 0.9537 | 0.9706 |
| Stem diameter/Stem diameter | 0.8441 | 0.8602 | 0.8367 | 0.8346 |

### 3D root traits distinguish genotypes in the test panel

A principal component analysis of 29 DIRT/2D traits suitable for maize root crowns and the 18 DIRT/3D root traits show both distinguishable clusters per genotype in the projection on the first and second principal component. In our test data set, the first two principal components explain 48.7% of the overall observed variance in DIRT/2D (Figure 9a), which compares to 51.9% explained variance in DIRT/3D (Figure 9b). Both, 2D and 3D root traits also distinguished genotypes by their normalized mean values (Figure 9cd). We found that no single trait classifies all genotypes in 2D and 3D. However, an ANOVA test revealed that the means of each pair of genotypes distinguishes in at least three traits for DIRT/2D and four traits for DIRT/3D (Supplementary Material 12). For example, in DIRT/3D genotype PA762 and B101 show a significant difference with traits such as nodal root diameter in the youngest whorl and lateral root diameter. However, B101 and PHG50 X PHG47 do not show separable mean values in the nodal root angles. We excluded the whorl distance between the 2^nd^ and 3^rd^ youngest whorl from the analysis because the distance was not detectable for some genotypes.

**Figure 9:**
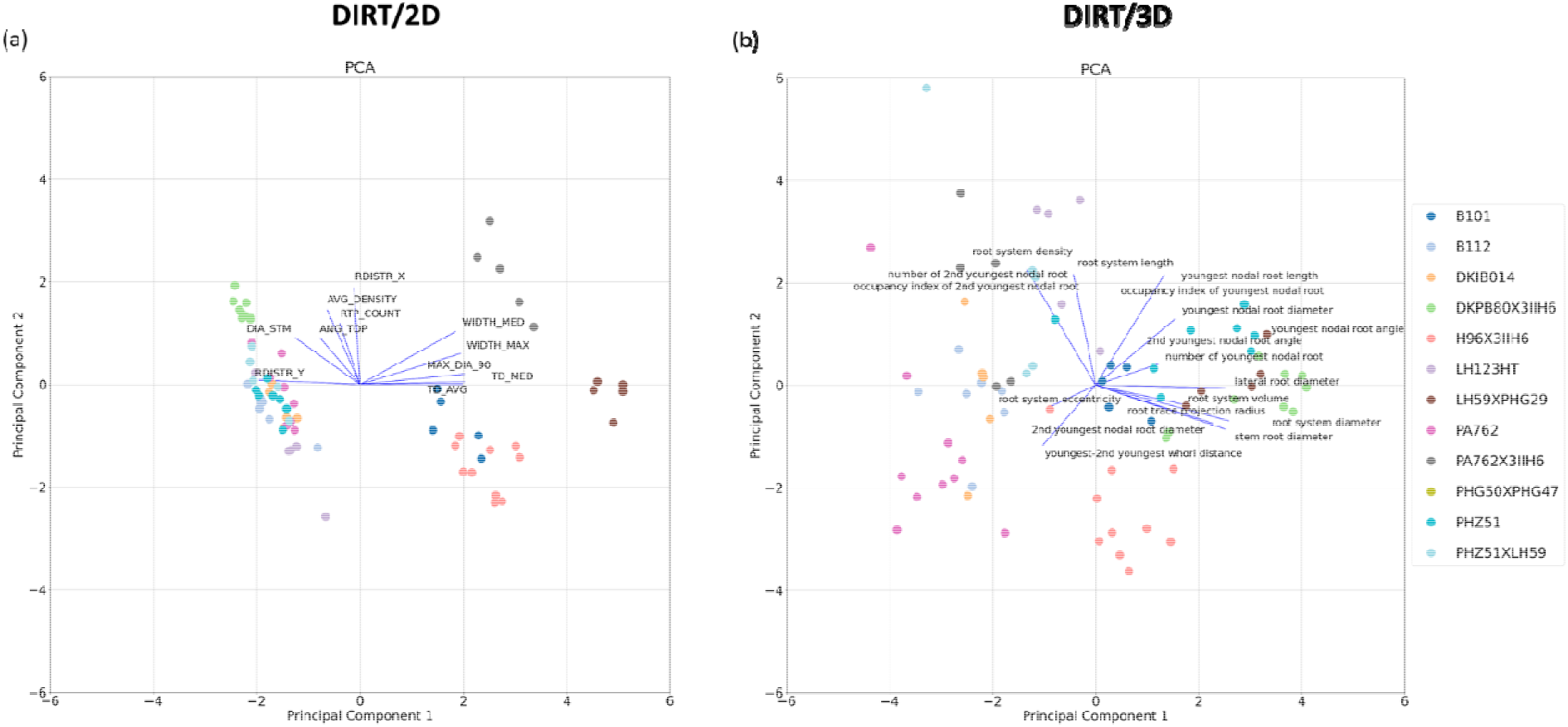
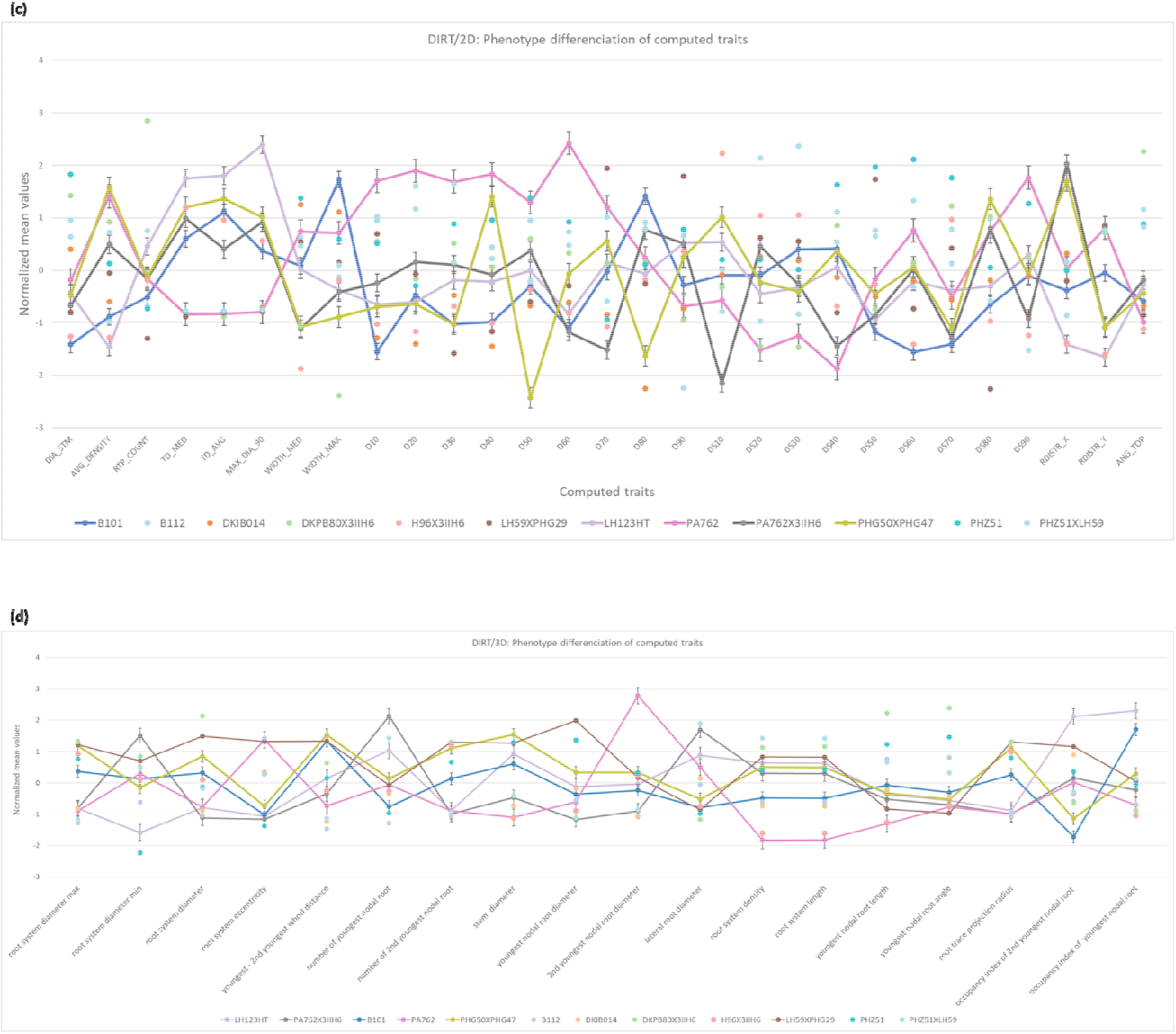
Genotype differentiation of 12 maize genotypes. (a) Principal component analysis (PCA) of DIRT/2D traits suitable for maize root crowns. (b) Improved maize root PCA using DIRT/3D traits. Colors correspond in (a) and (b) to genotypes and points to measured root crowns. We normalized all the mean trait values of computed 2D and 3D root traits from DIRT/2D (c) and DIRT/3D (d). Colored points denote the normalized mean values of the 16 root traits and error bars correspond to the standard error of the mean. The lines guide the reader visually to explore the phenotypic variation between genotypes of the test panel. For example, the mean of genotype PA762 and B101 distinguishes in nodal root diameter of the youngest root forming whorl and lateral root diameter in case of DIRT/3D; Genotype PHG50 X PHG47 and PA762 distinguish in the root projection radius. However, B101, PA762, and PHG50 X PHG47 do not show distinguishing mean values in the nodal root angle of the youngest whorl.

### Whole root descriptor distinguishes the unique spatial arrangement of individual roots for all genotypes

Figure 9c illustrates that D- and DS-values, which are samples of the D- and DS- curve, add strong discriminating power to DIRT/2D. Therefore, we introduce a 3D variation of the established D-curve for 2D images (Bucksch et al., 2014) as a whole root descriptor with improved differentiation capabilities (Figure 10). We compute the descriptor from the sequence of level set images derived from the reconstructed 3D root models. For each level set image, we compute the number of pixels that represent roots as a measure for the area. We found that the accumulation of root area across the level-set images is an intrinsic characteristic of each genotype (Supplementary Material 13). The descriptor is robust to outliers and measurement errors because it relies on the cumulative distribution function (Chun et al., 2000; Lee, 2001; Kyurkchiev, 2015). The 3D whole root descriptor distinguished the unique arrangement of individual roots for all 12 genotypes as a characteristic mean curve (Figure 10b). In comparison, the 2D whole root descriptor (Figure 10a) requires further downstream processing of the curve shape to achieve comparable distinction with derived descriptors like the DS-curve (Supplemental Material 13 and Figure 9b) (Bucksch et al., 2014). While whole root descriptors are powerful tools to capture the shape of a root crown or a branching structure in general, they also add value to commonly used analysis methods. In Figure 10c and d. we added D/DS-values and CDF values respectively to the PCA shown in Figure 9a and b. In both cases an improvement of the clustering is visible. For DIRT/2D the explained overall explained variance improved from 48.7% to 49.7%and for DIRT/3D the overall explained variance improved from 51.9% to 53.2%.

**Figure 10:**
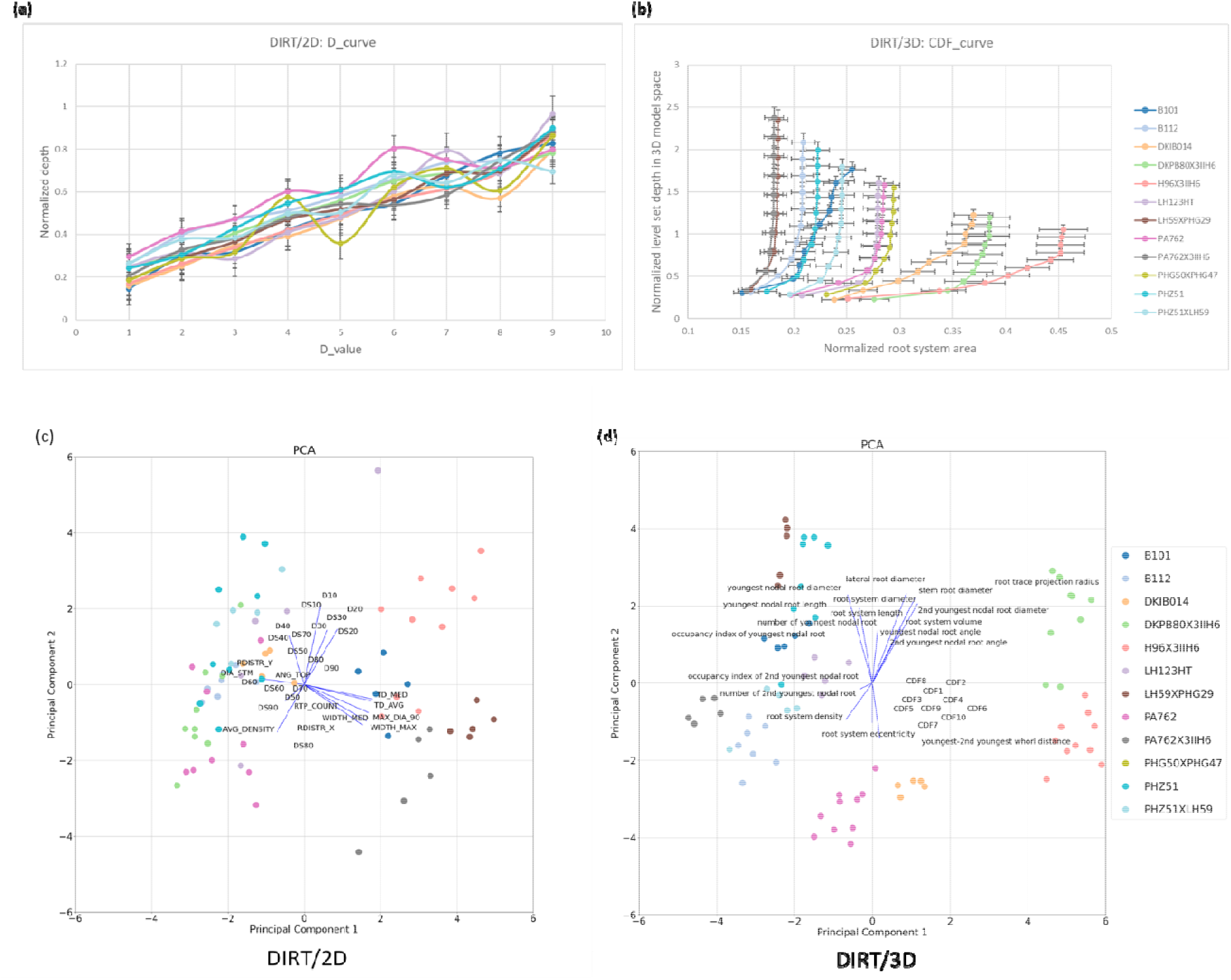
Comparison of 2D and 3D whole root descriptors of all 12 maize genotypes of the test panel. (a) Accumulating root width in 2D instead of accumulating volume in 3D results in differences in the curve shape. Visually wavy, straight and irregularly shaped curves can be distinguished. However, analyzing such shapes needs further downstream analysis of curve behavior such as with derived DS-curves (Supplemental Material 12). (b) The 3D descriptor encodes the spatial arrangement of individual roots within the root crown as a function of the excavation depth. We define the curve of the cumulative root crown area as the cumulative distribution function (CDF) of the area per level set for each genotype. The error bar denotes the standard error of the normalized root area. Each genotype associates with a characteristic CDF curve (colored coded). All genotypes distinguish visually from each other in their curve characteristics. (c) Sampling the D and DS curve at fractions of the excavated depth improves clustering of genotypes in the 2D PCA. (d) Adding the CDF curve traits to the PCA results in visually clear distinction of the genotypes.

## Discussion

The presented 3D system to measure traits in highly occluded root crowns is a significant advance in root phenotyping because it measures highly occluded traits such as whorl distances and number of nodal roots in dense maize root crowns. A main difference of the presented 3D system to other 3D systems, like X-Ray and MRI, is that no special infrastructure such as radiation protected rooms or specialist training for personal is required. Furthermore, the presented methods introduce a new 3D whole root descriptor for plant root architecture. For the test panel of twelve genotypes, a minimum of four traits distinguished all genotypes. In contrast, the whole root descriptor distinguished all genotypes with one mathematical expression. With these data, we have made significant progress on the unaccomplished goal of phenomics to measure the comprehensive appearance of a continuously reshaping phenotype (Houle et al., 2010).

From a validation point of view, the top angle (r^2^=0.87) and median root crown width (r^2^=0.88) published in the 2D DIRT system (Das et al., 2015) show equally good correlations with manual measurements as the comparable nodal root angle of the youngest whorl (r^2^=0.88) and root crown diameter (r^2^=0.89) on a maize data set. Certainly, comparable results could be expected for the nodal root angles of the second youngest whorl if occluding roots were mechanically removed from the root crown. Here we reproduced the previously reported 2D correlations to a manually measured ground truth. For comparable traits, we found that DIRT/3D achieved very similar and high correlations (r^2^>0.84) with the ground truth data for traits in both the DIRT/2D and DIRT/3D output. Broad-sense heritability also showed high values of H^2^>0.6 for the compared traits for DIRT/2D and DIRT/3D. However, the presented 3D system extends the availability of traits compared to DIRT/2D in two ways: Firstly, many occluded traits can be revealed without mechanical work and secondly, traits like nodal root number at specific nodes or root crown eccentricity can be estimated without additional manual data collection. Nevertheless, some algorithmic and technical challenges remain to exploit the utility of 3D root phenotyping for breeders fully. To date, standard calibration procedures for structure from motion scanners with multiple cameras are rare (Conte et al., 2018) and limit the achievable resolution to detect nodes within the maize root crown. Further research will focus on the details of the photogrammetric calibration of the 3D root scanner, which will allow for thinner cross-section slices during level set extraction. We believe that root models of higher resolution will enable the reconstruction of a more detailed root architecture to obtain measures of all nodes in the maize root crown.

We demonstrated the possibility to retrieve high geometric detail from field excavated root crowns. We even argue that it will be possible to obtain geometrically complete measurements in the sense of Euclid’s definitions in Elements I-IV and VI (Callahan and Casey, 2015). Local measurements of length, diameter, and angle are sufficient to reconstruct every solid 3D object if sampled at sufficiently high rates. Again, research on the calibration technique used for structure from motion scanners seems likely to be the limiting factor in achieving the needed resolution. Assembling the complete geometrical system of the root crown will allow us to describe the whole root crown and its spatial arrangements in one single mathematical construct. We presented a 3D whole root descriptor that is methodologically similar to the D- and DS-curve for 2D images (Bucksch et al., 2014). Here, we encoded root architecture as an aggregate of the extracted 3D root volume and could reliably distinguish the roots of different genotypes for a small diversity panel. While, our approach varies dependent on the spatial arrangement of roots, it does not encode the branching structure explicitly. An extension of the presented whole root descriptor would enable the quantification of morphological differences to understand the variation of architecture arrangements. Besides, the comparison between plant species with similar topological organization but different geometric growth such as dense first order laterals yet with varying patterns of curvature along the root, would be enabled.

The observed broad-sense heritability suggests a strong repeatability with *H^2^_mean_* > 0.6 for all traits except nodal root length of the youngest whorl. Repeatability, paired with near geometric completeness, indicates the presence of a local and global architecture control by genes. Local control relates to phenes that assemble the architectural phenome of roots as a set of mappable and locally measurable traits (Lynch and Brown, 2012). However, it is still an open question if a “global control phene” of root architecture phenes exists (Jiang et al., 2019) unless we can map whole root architectures that are geometrically complete. A necessary step towards answering this question from a mathematical point of view is to define a mathematical basis of locally controlled traits or phenes. Since phenes are often mappable to genes (Yablokov, 1986), a mathematically independent basis formed by phenes would open ways for the alternative hypothesis that roots have access to a spectrum of architectures that acclimatize to their micro- and macro-environments via their species-specific phenes.

Our system will improve its capabilities with more 3D data becoming available that enables deep learning approaches on the 3D root data. For example, the distinction of brace and crown roots instead of root forming whorls depends on the detection of pigmentation on the aerial nodal roots. As more 3D datasets become available, deep learning networks can be trained to differentiate between crown and brace roots based on color and texture differences that vary with environment and genotype. This current data limitation demands careful planning of the excavation protocol and computing setup to distinguish brace and crown whorls e.g. as youngest or 2^nd^ youngest whorl. Therefore, we currently resolve five to six root forming whorls dependent on the genotype. The number of detected whorls is a resolution limited estimate assuming two whorls formed during early root development. These two early whorls cannot be distinguished given the resolution limit of 2 mm whorl distance. However, resolution limits likely also apply to any other technology because the oldest whorls often remain even visually indistinguishable.

## Conclusion

Our 3D phenotyping system is an optical system to handle highly occluded and mature root crowns collected in the field. It is worth noting that the time required to collect the imaging data is around five minutes, which is similar than an X-Ray scan at a comparable resolution (Jiang et al., 2019). Unlike many root phenotyping methods developed in controlled environments, our system measures maize roots grown under field conditions. Our results demonstrate that some traits such as root width achieve comparably good correlations with manually measured ground truths and broad sense heritability values with 2D and 3D phenotyping systems. However, we also demonstrated the added value of DIRT/3D is to reliably compute traits that are inaccessible in 2D images. In particular, dense root crowns like maize suffer from the unavailability of detailed traits in 2D. Such traits include whorl distances, the number and angles of nodal roots forming the youngest or 2^nd^ youngest whorl. We validated our system for the root trait classes of number, angle, diameter, and length. Validation results demonstrate the reliability of our system with correlations of r^2^ > 0.84 for all traits and P < 0.001. From our analysis, we concluded that DIRT/3D could extract 3D root traits accurately at the individual and crown levels.

We also demonstrated that whole root descriptors improve the capabilities for analyzing root phenotypes. Whole root descriptors quantify the overall shape of the root an allow for downstream shape analysis of different root shapes. A second added value of whole root descriptors is to strengthen cluster differentiation of PCA analyses. In our example we found strong evidence that whole root descriptors improve genotype differentiation with PCA analysis. The improvement is visible in the projection of the first two principal components. Augmenting PCA analysis with whole root descriptors resulted in explained variance of 49.7% in DIRT/2D and 53.2% in DIRT/3D.

Both, our software and hardware design are an open and inexpensive 3D root phenotyping solution. At time of publication, the complete system was developed for about $6000, which includes labor costs to produce the frame and high-end cameras. We currently explore options to build the complete 3D system for about $1500-2000 using cheaper cameras and other means to produce the rotation stand. Our open-source software is available to the whole plant science community on GitHub and can be deployed within a platform-agnostic Singularity/Docker container to be executed independently of the operating system (Supplementary Material 14) (https://github.com/Computational-Plant-Science). The use of Singularity/Docker containers will allow for integration with cyber-infrastructures such as CyVerse. These containers can run on any high-performance computing system that has the Singularity environment installed. The scanner design is part of the publication (Supplementary Material 3) and can be reproduced, scaled and further developed by everyone.

The presented 3D system requires only one user interaction to place the root crown in the scanner. Placing the root in the scanner could be replaced by a robot in future. Hence, we see our system as the first milestone towards automated root trait measurements in the field. Our belief stems from ongoing developments in agricultural robotics that will excavate field roots “on-the-go” (Shi et al., 2019) in the foreseeable future. In that way, our system supports breeders and root biologists in the development of crops with increased water uptake, more efficient nitrogen capture and improved sequestration of atmospheric carbon to mitigate the adverse effects of climate change without compromising on yield gains.

## Material and Methods

### Plant material

Plants were grown at The Pennsylvania State University’s Russell E. Larson Agricultural Research Center (40° 42’40.915” N, 77°, 57’11.120’’W) which has a Hagerstown silt loam soil (fine, mixed, semi-active, mesic Typic Hapludalf). Fields received fertilization with 190 kg nitrogen ha^−1^ applied as urea (46-0-0). The sites had drip irrigation. The field management supplemented nutrients other than nitrogen, and applied pest management as needed. We planted seeds using hand jab planters in rows with 76 cm row spacing, 91 cm alleys, 23 cm plant spacing, 4.6 m plot length with 3.7 m planted, or ~56,800 plants ha^−1^. We grew plants in three-row plots and sampled only the middle row. Planting occurred on June 5, 2018, and sampling on August 25, 2018, 81 days after planting. Two fields provided 1ha of space for four replicates.

Twelve genotypes were selected based on previous knowledge of their architectural variation and sampling of a larger set of genotypes. The twelve genotypes included six inbred lines (B101, B112, DKIB014, LH123HT, Pa762, PHZ51) and six hybrid lines (DKPB80 x 3IIH6, H96 x 3IIH6, LH59 x PHG29, Pa762 x 3IIH6, PHG50 x PHG47, PHZ51 x LH59). These genotypes represent the extremes of dense vs. sparse, large vs. small, and maximum and minimum number of whorls selected from a full diversity panel. The lab of Shawn Kaeppler at the University of Wisconsin provided the seeds. We selected ten representative plants for five of the genotypes (B112, Pa762, PHZ51, DKPB80 x 3IIH6, H96 x 3IIH6), and five representative plants of the remaining seven genotypes. Sampling followed the shovelomics protocol (Trachsel et al., 2011), which minimizes variation by selecting similar representative architectures. Shoots were removed above all root-producing nodes. We air-dried the roots on a greenhouse bench and then transported the roots to the lab for imaging.

### Obtaining the ground truth for root trait validation

Each root crown was fixed on a board. We used a ruler to measure the length and diameter of the root crown. A second diameter was measured orthogonal to the board plane to determine the eccentricity of the root crown. We used a protractor to measure the nodal root angles to the horizontal from four sides. The average angle of the four sides was taken to represents the rooting angle for nodal roots forming respective whorls. To measure root diameters and whorl distances, we used a steel vernier caliper with a graduation of 0.02mm. The came caliper was used to measure maximum root crown diameters. The dry weight of the root crowns was weighed with an ADAM Core Portable Compact Balance CQT 202 (readability: 0.01g, linearity: 0.02g).

### 3D root scanner

We designed a 3D root scanner (Figure 2a) to capture images for 3D reconstruction of the root (Supplementary Material 3). A stepper motor (Nema 34 CNC High Torque Stepper Motor 13Nm with Digital Stepper Driver DM860I, Figure 2b) rotates a curved metal frame with ten low cost and highly versatile imaging cameras (Image Source DFK 27BUJ003 USB 3.0, 6mm focal length and TCSL 0618 5MP lens) around the clamped root crown in a central fixture (Figure 2c). From the stepper motor, we chose 12800 micro-step resolutions to rotate in 1-degree steps (Figure 2b). The cameras ship with the 1/2.3” Aptina CMOS MT9J003 sensor and can achieve high image resolution at 3,856×2,764 (10.7 MP) up to 7 fps. We drilled 21 equidistant holes into the curved frame to provide a flexible arrangement of each camera. A rail track along the curved frame allows for fine adjustment of the camera tilt and pan direction without compromising stability (Figure 2d). Cameras are then arranged along the curved frame to achieve a sampling of bigger and smaller root morphology that satisfy the Nyquist theorem to prevent aliasing (Liu et al., 2009). In the case of maize roots, more cameras are concentrated to image the root crown with high amounts of small occluded roots. Only two cameras cover the stem part of the because the surface area of the stem part usually has minimal to no conclusion, which guarantees good 3D reconstruction results.

A computing cluster of ten Raspberry Pi 3 B+ synchronizes the image capture of the ten cameras using a server-client design (Supplementary Material 15). The synchronized cameras of our 3D root scanner capture approximately 2000 images per maize root in about 5 min. The newly developed controller software on the Raspberry Pi computing cluster synchronizes the camera’s image capture and the stepper motor movement. Once the stepper motor receives the “start move” signal via the server unit, it moves all the cameras into their designated positions. Then, all ten cameras capture images simultaneously. Each Raspberry Pi stores the image initially on a sim card. During the image capturing process, the stepper motor stands still and waits for the next “start move” signal. The image data of all Raspberry Pi’s automatically transfers to the CyVerse Data Store (Goff et al., 2011; Merchant et al., 2016). Only the server unit stores information about the CyVerse user account. It uses the iRods protocol (Ward et al., 2009) to transfer the images from each client unit to the CyVerse Data Store. In the following, the 3D reconstruction uses the image data in the online storage to generate the 3D point cloud of the root crown. Alternatively, the image data can be transferred manually to computers within the same WiFi network.

### Automatic reconstruction of the 3D root model with structure-from-motion

#### Fast Fourier Transform (FFT) detects blurry images

The structure from motion (SfM) method requires detected feature points to be visible in several camera views. However, pose inaccuracy mechanically inferred by the scanning device or false feature matching may lead to incomplete reconstructions (Zheng and Wu, 2015). As a result, not all feature points are triangulated to generate 3D points. In our case, a small number of images acquired with the 3D scanner appear dark or blurred because of delayed image capture, frequency of surrounding light sources or vibrations of the scanner (see Supplementary Material 16 for an example). We detect blurred images using Fast Fourier Transform (FFT) to transform the image into the frequency domain. The absence or low number of high frequencies compared to the majority of images indicates a blurred image. Removing blurred images results in higher confidence for feature matches and therefore, improves model reconstruction quality and point density in SfM approaches.

#### Illumination adjustment and content-based segmentation to remove redundant information

We use standard deviation and a luminance-weighted gray world algorithm (Lam, 2005) to adjust and normalize illumination across all captured images. The root is automatically separated from the background using a newly developed content-based segmentation method (Supplementary Material 17). The method analyzes and compares color-space differences across all normalized images and omits the redundant information of the background. Overall, the size of the image data reduces to 30-50% of the original size. In later steps of the pipeline, the segmentation decreases the number of false feature matches during the 3D reconstruction process as well as the amount of data transmitted to online storage. The method is fully automatic and parameter-free and uses parallel processing if available.

#### Improved feature matching to reduce computation time and improve 3D point cloud resolution

Given the images of segmented roots, we chose the Visual Structure From Motion method (Wu, 2011) as a basis to develop 3D reconstruction software for roots. The computationally most expensive aspect of structure-from-motion algorithms is the feature matching between image pairs. The amount and accuracy of the feature matching determines the quality and resolution of the resulting 3D root model. In the original version, Visual Structure from Motion performs a full pairwise image matching to build a feature space across all possible image pairs. For example, the number of permutations P calculates for r images out of a set of n total images with the following formula:

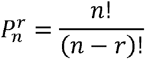

However, the computation of feature matches generates a large amount of false feature correspondences in the dense root data. We found that image pairs that are not adjacent in the spherical scanner space are particularly prone to incorrect matching (Supplementary Material 18). We observed that the false feature matches occur predominantly between the dense and thin roots of the root crown. Therefore, we optimized the feature matching process to be suitable for dense root architectures.

The optimization in our algorithm generates a matching pair list inside a specified sliding window (Supplementary Material 19). Sliding of the window allows for robust matching among all permutations of image pairs. For example, given an image set captured around the individual root in the 1-degree interval (360 images in total), we set the sliding window size as 10% of the image size. The widow size was found experimentally and is the optimum for the 1-degree interval setting of the scanner. The total number of permutations of image pairs needed for feature matching is 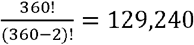 according to the formula above. For an image of size 1000 × 1000, we set the sliding window size as 100×100, the number of permutations of image pairs needed will be reduced to 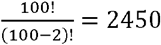. In that way, we need to compute only 1.89% of all permutations of image pairs.

As a next step, we utilize the RANSAC (random sample consensus) method to detect and remove the falsely matched pairs. The RANSAC results usually contain only highly distinctive features to track between consecutive images. Given the locations of multiple matched feature pairs in two or more images, we can produce an estimation of the positions, orientations of cameras, and the coordinates of the features in a single step using bundle adjustment (Wu et al., 2011).

#### Computing root traits from 3D models

We adopted a top-down level set scan of the 3D root model to compute 3D root traits (Supplementary Material 20). This scanning process generates a thin vertical 2D slice or level set image given a fixed plane. We use a phase-based frame interpolation technique from video processing to smooth the image sequence. We developed a method to extract individual roots in each level set image using the active contour snake model. Then we use the watershed segmentation to segment the overlapping roots. Given a smoothed and segmented level set image sequence, we used a combination of Kalman filters and the Hungarian algorithm (Sahbani and Adiprawita, 2016) to track all individual roots, and build an embedded graph of the geometry of resolved nodal roots at their respective nodes. This embedded graph forms the basis to compute all 18 root architecture traits. Nodal root traits are directly derived from the embedded graph. In each level set image, we compute the lateral root diameter as an average of the circular projections of point cloud points that are not identified as nodal roots emerging from a resolved node. For each level-set image, we compute the area covered by nodal roots of resolved nodes. This area increases whenever nodal roots emerge from a whorl and stays almost constant between whorls. If summarized as a cumulative function of area (see Supplemental Material 13) the starting location in the level set image stack corresponds to the starting point of a plateau in the cumulative function.

#### Statistical Analyses and used software

All statistics used python 3.7 and the modules NumPy 1.16 and SciPy 1.2.1 (Oliphant, 2007). Figures 7 and 10 used matplotlib 3.2.1 (Hunter, 2007) for visualization of the statistics. Figures 8 and 9 used Microsoft Excel Version 16.34 to visualize trait and heritability data. Raw data are available in (Supplementary Material 12).

DIRT\2D traits were computed with the online platform available at http://dirt.cyverse.org on May, 4^th^ 2021.

## ACKNOWLEDGMENTS

The research was supported by the USDOE ARPA-E ROOTS Award Number DE-AR0000821 to A.B. and J.P.L.. The work was supported in part by the NSF CAREER Award No. *1845760* to A.B. Any Opinions, findings, and conclusions or recommendations expressed in this material are those of the author(s) and do not necessarily reflect those of the National Science Foundation. This work used the Extreme Science and Engineering Discovery Environment (XSEDE) resource Stampede2 at the Texas Advanced Computing Center through allocation TG-BIO160088. This material is partly based upon work supported by the National Science Foundation under Award Numbers DBI-0735191, DBI-1265383, and DBI-1743442. URL: www.cyverse.org.

## CONTRIBUTIONS

S.L. wrote the manuscript, designed and implemented algorithms, designed and built hardware, performed data analysis and contributed to the experimental design. C.B.S. designed and built hardware. J.P.L. contributed to writing of the manuscript and the project idea, conceived experimental design. M.H. contributed to writing the manuscript, performed experiments and collected data. A.B. conceived the project idea, designed hardware, contributed to the data analysis, wrote manuscript and designed algorithms.

## Notes

### Competing Interest Statement

The authors have declared no competing interest.

